# Performance in a novel environment subject to ghost competition

**DOI:** 10.1101/515833

**Authors:** Karen Bisschop, Frederik Mortier, Dries Bonte, Rampal S. Etienne

**Author notes:** Joint last authorship (Dries Bonte, Rampal S. Etienne). Corresponding author: Karen Bisschop.

## Abstract

**Background:** A central tenet of the evolutionary theory of communities is that competition impacts evolutionary processes such as local adaptation. Species in a community exert a selection pressure on other species and may drive them to extinction. We know, however, very little about the influence of unsuccessful or ghost species on the evolutionary dynamics within the community.

**Methods:** Here, we studied the long-term influence of a ghost competitor on the performance of a more successful species using experimental evolution. We transferred the spider mite *Tetranychus urticae* onto a novel host plant under initial presence or absence of a competing species, the congeneric mite *T. ludeni*.

**Results:** The latter species unintentionally went extinct soon after the start of the experiment, but we nevertheless completed the experiment and found that the initial density of this ghost competitor positively affected the performance (i.e. fecundity) of the more successful species.

This effect on *T. urticae even* lasted for at least 25 generations.

**Discussion:** Our study supports the hypothesis that early experienced selection pressures can exert a persistent evolutionary signal on species’ performance in novel environments.

## INTRODUCTION

Species are facing a continuously changing world that they can possibly cope with in various ways, such as through phenotypic plasticity or by tracking their favoured habitat. If these solutions are not possible, evolutionary rescue by genetic adaptation may eventually allow persistence (Lindsey et al., 2013). One factor influencing local adaptation is competition, which can occur among con- or heterospecifics. Niche overlap is usually larger within than among species (Bolnick, 2001; Svanbäck & Bolnick, 2007); therefore, it is expected that competition between conspecifics has a larger impact on evolution than competition at the interspecific level.

Interspecific competition is also known to influence local adaptation, but the effect is still largely unpredictable (Rice & Knapp, 2008; Alzate et al., 2017; Zhao et al., 2018). First, heterospecific competitors might modify the selection pressure exerted by the abiotic environment, enhancing or limiting genetic adaptation to the novel environment (Osmond & de Mazancourt, 2013). Classical examples of enhanced genetic adaptation are seen in adaptive radiations of three-spined sticklebacks or fast character displacements in Darwin finches or Myzomelid honeyeaters (Diamond et al., 1989; Schluter, 1994; Reznick & Ghalambor, 2001). Previously, we found that additional selection pressure exerted by a congeneric species facilitated adaptation of the focal species to a novel environment under high dispersal from a maladapted ancestral population (Alzate et al., 2017). Adaptation to the novel environment can also be reduced by interspecific competition when there is, for instance, a trade-off between traits responsible for adaptation to the competing species and to the novel environment (Siepielski et al., 2016).

Furthermore, interspecific competition can create new niches or change the current environment for species to adapt to. Species may use waste products or adapt to plants with modified defences caused by coexisting individuals (Sarmento et al., 2011; Lawrence et al., 2012). These new niches will subsequently create opportunities for adaptive shifts to novel environmental conditions. This illustrates that competition and facilitation can jointly shape evolution, making it difficult to study the consequences of interspecific competition alone.

As a last scenario, interspecific competition can hinder the process of local adaptation by restricting resource availability and hence decrease effective population size. The resulting increased probabilities of genetic drift will then decrease the evolutionary potential and hence the chance of local adaptation (Lawrence et al., 2012; Osmond & de Mazancourt, 2013; Zhao et al., 2018).

While inferior competitors are expected to eventually go extinct, they may coexist with the more successful competitors for many generations (Holmes & Wilson, 1998; Lankau, 2011). These early and non-persisting interactions may leave a strong signature on the future community dynamics (Law & Daniel Morton, 1996; Miller, TerHorst & Burns, 2009; Mallon et al., 2018), because they have the possibility to induce large habitat modifications or evolutionary changes in the more successful species. Historical contingency (i.e. the influence of the arrival time of a certain species in a community; Fukami 2015) in terms of limitations imposed by so-called ghost species (Hawkes & Keitt, 2015), may thus have a strong impact on the eco-evolutionary trajectories of populations and communities, in the same way as successful species do (Fukami, 2015). The role of competition intensity of an inferior species prior to its extinction on the ecological and evolutionary dynamics of persisting species is still largely unknown, however. Here we present results from an evolutionary experiment with two related spider mite species adapting to a novel host. Both species were placed alone or together on a new host plant and we wanted to verify how this interspecific competition affects local adaptation. The two competitors were supposed to be competitively similar, but the experiment demonstrated that this was not the case: both species could only temporarily coexist. This provided us the unique opportunity to investigate the influence of ghost competition on adaptation. More precisely, we explored whether we could detect long-term evolutionary effects on performance (measured as fecundity) due to differences in initial selection pressures caused by this ghost competitor. We chose the average of the initial population size of the unsuccessful species during the first month of coexistence as an indication for the initial selection pressure. These differences in population sizes arose naturally and can be attributed to selection, as well as drift and founder effects. *T. ludeni* showed a lower fecundity than *T. urticae* on bean and cucumber in the control populations, which may explain their rapid extinction. Still, the ghost species *T. ludeni* showed an effect on the surviving species *T. urticae*, because the eventually achieved strength of adaptation of *T. urticae* increased with the initial density of *T. ludeni*. We therefore suggest that ghost competition is an underestimated process for adaptation and may lead to differences in long-term local adaptation.

## MATERIAL AND METHODS

### Study species

We used two species of the family Tetranychidae (Acari, Arachnida): *Tetranychus urticae* Koch, 1836, and *T. ludeni* Zacher, 1913. These herbivorous mite species are highly suitable for evolutionary experiments due to their small body sizes, their possibility to maintain large populations in the lab, and short generation times (Zhang, 2003).

For this study, we used inbred populations of *T. urticae* from Bitume and colleagues (2013). Each population originated from two adult females from the LS-VL line (Van Leeuwen, Stillatus & Tirry, 2004) and was afterwards kept at low population densities. The LS-VL line was collected from roses in October 2000 (Ghent, Belgium). After this initial collection, all populations were maintained on bean plants (*Phaseolus vulgaris,* Prelude).

Two populations of *T. ludeni* were used: the Tl Alval (Lisbon, Portugal) and Tl CVM (Lourinhã, Portugal). Both populations were sampled early autumn 2013 from common morning-glory and afterwards maintained on bean plants (*P. vulgaris,* Prelude). The founder populations were 160 and 300 individuals for Tl Alval and Tl CVM respectively. Our evolutionary experiment started in September 2015, implying that *T. urticae* and *T. ludeni* had been under laboratory conditions for about fifteen and two years, respectively.

For this study, we chose to subject the inbred lines of *T. urticae* (Bitume et al., 2013) to further inbreeding by mother-son mating for one more generation prior to the experiments. This resulted in the creation of 13 isofemale lines. It may sound counterintuitive to use inbred populations for an evolutionary experiment that mainly uses standing genetic variation (note that no spider mites were added during our experiment), but in this way we could generate genetically similar replicates and control for putative initial drift effects by differences in starting genetic variation. We deemed this more important than potential inbreeding effects, because no effects of inbreeding on genetic trait variation were found in these and other lines (Van Petegem *et al*., 2018; Bonte unpub. results). We additionally created six isofemale lines for *T. ludeni* (coming from Tl Alval and Tl CVM). We wanted to create 13 lines for this species as well, but were unsuccessful due to low fertility or early mortality. The stock Tl Alval and Tl CVM populations were placed on bean plants (four two-weeks-old plants) and are from here on referred to as the control population of *T. ludeni.* A control population of *T. urticae* was also created from the created 13 isofemale lines (four mites per line) on bean plants (four two-weeks-old plants). All populations were kept in a climate-controlled room (25°C – 30°C, 16:8 L:D).

### Experimental set-up

At the beginning of the actual experiment, the isofemale lines were placed on novel host islands (two three-weeks-old cucumber plants, *Cucumis sativus* Tanja, per island) with or without heterospecifics. After the first week, two fresh three-weeks-old cucumber plants were added to create the island size of the experiment. Afterwards, the islands were weekly refreshed by replacing the two oldest plants with two new three-weeks-old cucumber plants. In this way, sufficient time was provided for a generation of spider mites to develop on the new plants, while allowing the population to move toward the fresh leaves. Hence, while the removed old plants may have contained mites or unhatched eggs, we chose for this refreshment procedure to maintain natural movement dynamics. It is for instance known that especially young fertilised females disperse more (Li & Margolies, 1993) and dispersive individuals may differ in their body condition or performance compared to sedentary individuals (Bonte et al., 2014; Dahirel et al., 2019). This refreshment procedure may have caused an extra competitive pressure if one species was more dispersive or delayed its dispersal for avoiding competition, but we preferred to design the experiment in a way that it resembled more the actual life strategy of spider mites (colonisation with few founders followed by rapid growth).

The novel host islands were placed in boxes with yellow sticky paper (Pherobank) at the bottom and Vaseline at the walls to avoid contamination between islands; this method is known to work from previous research (Alzate et al., 2017; Alzate, Etienne & Bonte, 2019; Bisschop et al., 2019). Eight islands (or replicates) received both *T. urticae* and *T. ludeni.* Eight islands (or replicates) received only *T. urticae* and another eight islands (or replicates) received only *T. ludeni*. Each island started with the same total population size and as similar as possible gene pool. The group with both spider mite species received 26 adult females of *T. urticae* (two from each of the isofemale lines) and 26 adult females of *T. ludeni* (resulting in 52 adult females). Twelve *T. ludeni* females per island came from the six isofemale lines and were supplemented with 14 mites from its stock population, because of the lack of success for creating more isofemale lines. The use of the outbred stock population of *T. ludeni* to supplement the populations provided an unanticipated opportunity. In this way, we could benefit from the larger initial genetic variation of *T. ludeni* among replicates and hence differences in initial selection pressures on *T. urticae.* The group with only *T. urticae* received four adult females from each of the thirteen isofemale lines (resulting in 52 adult females). The last group with only *T. ludeni* received four adult females from the six isofemale lines and was supplemented with 28 females from its stock population. We started with a rather low population size to make it biologically relevant as natural populations usually colonise plants at small population sizes. All adult female mites were equally distributed over the plants. In an ideal situation, the initial population sizes per species, the total population density, and the island size should be kept equal, but this is of course impossible. We chose for the same total population size and no differences among island sizes, as it is known that differences in densities change both the intra- and interspecific competitive pressure and that an increase in island size would change the adaptive potential of the treatment (Alzate, Etienne & Bonte, 2019). We acknowledge that this necessity of differences in initial population sizes might increase drift effects.

The total experiment lasted for ten months, which is about 25 generations and long enough to detect local adaptation (Gould, 1979; Fry, 1989; Magalhães et al., 2007, 2009; Bonte et al., 2010). For logistical reasons the experiment was performed in two blocks with one month difference, each block consisted of four replicates per treatment.

### Measurements

Every two weeks, the density of the spider mites in the evolutionary experiment was measured by counting adult females on a square of 1 x 1 cm²; the first counting was done after two weeks. The location of the square was right next to the stalk of the highest, fully grown leaf of the two newest plants of each island. Both the abaxial as well as the adaxial side were measured and summed for a total overview. The location on the leaf was chosen to standardise the measurements in time and make them comparable. The populations of *T. ludeni* under competition with *T. urticae* went extinct after about two months. To get an impression of its competitive pressure on the more successful *T. urticae* populations while it was still present, we used the mean population density of the first month of *T. ludeni*.

Fecundity tests for the control populations on bean and for the experimental cucumber populations were performed every two months to determine the level of adaptation. As the experimental populations of *T. ludeni* went extinct under competition, we obviously only have results from fecundity tests on the control population of *T. ludeni.* We chose fecundity as proxy of adaptation because previous research confirmed it to be the best predictor of adaptation compared to survival or development (Magalhães et al., 2007; Alzate et al., 2017; Alzate, Etienne & Bonte, 2019). Five adult females were sampled from each island and separately placed on a bean leaf disc (17 × 27 mm^2^) for two generations of common-garden to standardise juvenile and maternal effects (Magalhães et al., 2011; Kawecki et al., 2012). Bean discs were chosen because this is a very suitable host plant and will not cause a change in allele frequencies of the evolved lines (Magalhães et al., 2011). These leaf discs were placed in a petri dish on wet cotton wool and surrounded with paper strip borders. Then, the fecundity of two quiescent deutonymph females that originated from the same common-garden replicate was tested. One female was put on a bean leaf and one on a cucumber leaf (same set-up as for common garden) in a climate cabinet of 30°C under 16:8 L:D. Fecundity (number of eggs laid after six days) was measured based on daily pictures taken. Females that drowned in the cotton before the sixth day were excluded from the analysis (this was 13.5% for the populations of *T. urticae* without *T. ludeni*, 15% for the populations of *T. urticae* with the ghost competitor, and 10.5% for the populations of *T. urticae* in the control treatment maintained on bean). The cucumber plants for the fecundity test after four months did not grow for one of our two experimental blocks, so we were not able to test fecundity at that time point. In total, the fecundity was measured for 974 females (exact sample sizes per treatments are given in the electronic supplementary material Table S1).

### Statistical analysis

We used general linear mixed models (GLMMs) with Negative Binomial distribution with log link to account for overdispersion of the data. The variance was determined as µ * (1 + µ/*k*) in which µ is the mean and *k* is the overdispersion parameter (standard negative binomial parametrisation).

#### The dynamics and performance of the ghost competitor

We first studied the performance of the control populations that had been maintained on bean of both species on bean and cucumber, using Generalized Linear Mixed Models (GLMM). The dependent variable in the maximal model was fecundity (number of eggs after six days) and the explanatory categorical variables were the plant species (bean or cucumber) and the mite species (*T. urticae* or *T. ludeni*). Model selection was based on the lowest AICc and a Wald χ^2^ test was performed on the maximal model to check the reliability of the model selection. Pairwise comparisons were adjusted for multiple comparisons with Tukey’s method.

#### Signature of the ghost competitor on performance of T. urticae

We investigated the impact of the density of *T. ludeni* and of *T. urticae* at the onset of the experiment (i.e., mean density during the first month) on the fecundity of *T. urticae* on its initial and novel host plant. The explanatory variables in the maximal model were time (as categorical variable; 2, 4, 6, 8 or 10 months), the density of *T. ludeni* (continuous variable), the density of *T. urticae* (continuous variable), and the interaction between densities of both species and time. In this way, we aimed to determine whether it was the own density or the density of the ghost competitor that affected performance of *T urticae*. We compared this with an additional model with the total density (summing the density of *T. ludeni* and *T. urticae*), time, and their interaction to find out whether fecundity was affected by the species’ individual densities or just the total density. The random effects for all models were the island or replicate nested within the experimental blocks. However, we ran into convergence problems with the results from the assessment on cucumber as the random effect variance was estimated to be zero (Magnusson et al., 2018). As a consequence, we only used replicates as random variable. We chose a categorical variable for time instead of a continuous one, because differences in quality of leaves at the different measurements were likely and we did not want to assume a linear response of adaptation. Model selection was based on the lowest AICc and an additional Wald χ^2^ test was performed on the maximal model. Pairwise comparisons for the slopes and means were adjusted for multiple comparisons with the Tukey’s method.

We furthermore used the density assessed through time of the different spider mite populations to investigate differences in demography between the species with or without competitor. We divided the density through time in the first two months, when *T. ludeni* was still present under competition, and the last eight months. The dependent variable in the maximal model was the density (number of adult female mites per cm^2^) and the explanatory variables the treatment (*T. urticae* with and without competitor, and *T. ludeni* with and without competitor), time (continuous variable; linear for the first two months, second-degree polynomial for the last eight months) and their interaction. The random effects were the different islands or replicates within their experimental block. Model selection, Wald χ2 test, and pairwise comparisons were performed as explained above.

#### Performance without interspecific competitor

Because we were interested in the magnitude of the differences in performance due to the presence of *T. ludeni*, we did a further analysis including also the control population of *T. urticae* on bean and the populations of *T. urticae* on cucumber without *T. ludeni*. We investigated the fecundity in function of the three different treatments (control of *T. urticae* on bean, the populations of *T. urticae* on cucumber without *T. ludeni*, and those with *T. ludeni*) and time (categorical variable; 2, 4, 6, 8, or 10 months), and their interaction. The islands were treated as random effects and were nested within the two experimental blocks. Model selection, Wald χ^2^ test, and pairwise comparisons were performed as explained above.

The estimates provided in the tables are the raw and untransformed estimates for the fixed effects of the final models (negative binomial distribution). All analyses were performed in R (version 3.6.0) with glmmTMB version 0.2.3 (Brooks et al., 2017), MuMIn version 1.43.6 (Barton, 2019), emmeans version 1.3.5.1 (Lenth, 2019), and fitdistrplus version 1.0-14 (Delignette-Muller, 2015).

## RESULTS

### The dynamics and performance of the ghost competitor

In the competition treatments *T. ludeni* went extinct after about two months. While *T. ludeni* was able to maintain a population on cucumber in the absence of a competing species, it reached a significantly lower density than *T. urticae* (*t* ratio = 8.535 and *p* < 0.0001 for *T. urticae* under ghost competition; *t* ratio = 9.168 and *p* < 0.0001 for *T. urticae* without competitor). This suggests that the host plant itself is not a problem for *T. ludeni*, but that mainly the presence of *T. urticae* hindered the survival of the population (Fig. 1a, electronic supplementary material Table S2-S4).

**Figure 1:**
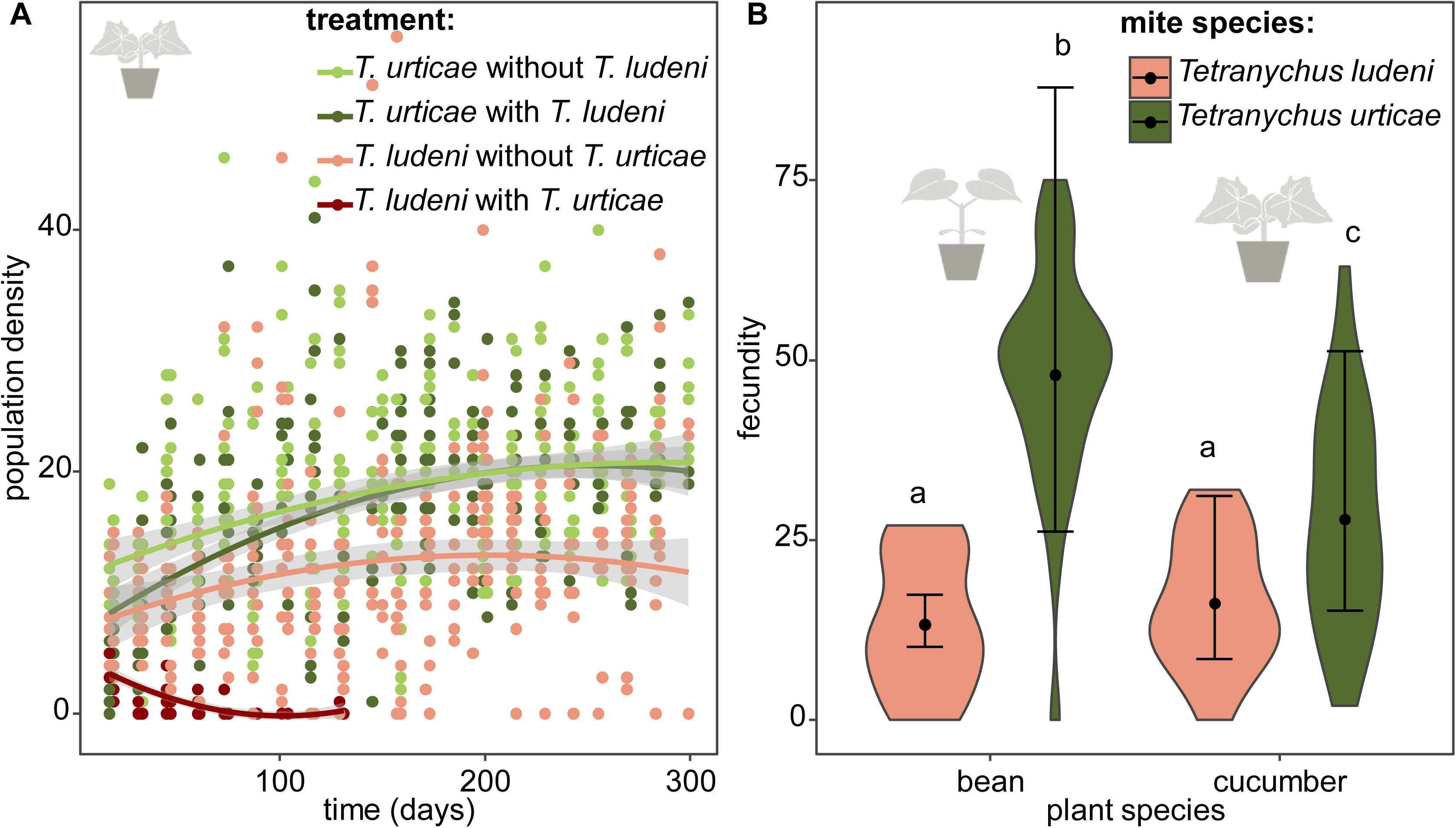
a) Overview of the population density for the different treatments. Population density of *T. urticae* (green dots) and *T. ludeni* (red dots) measured as the sum of the abaxial and adaxial density (number of adult females/cm^2^) per island through time. The lighter colours correspond to the populations in absence of the competing species and the darker to the treatment where both species are present. The grey zone shows the 95% confidence interval. **b)** Comparison of control populations of *Tetranychus ludeni* and *T. urticae* on bean and cucumber. The fecundity of *T. ludeni* is significantly lower than *T. urticae*, on both bean and cucumber. The violin plots show the observed data, and the points and lines show the mean model estimates and their 95% confidence interval, respectively.

The fecundity tests on the initial and novel host plant with mites from the control populations of *T. urticae* and *T. ludeni* (which had been maintained on bean plants and had never been on cucumber before) showed that *T. urticae* had a significantly lower fecundity on cucumber than on bean (*t* ratio = -3.629 and *p* = 0.0025), and that *T. ludeni* laid significantly fewer eggs on both bean (*t* ratio = -7.463 and *p* < 0.0001) and cucumber (*t* ratio = -6.177 and *p* < 0.0001) than *T. urticae*. There was no difference in the performance of *T. ludeni* on bean or cucumber (*t* ratio = - 1.012 and *p* = 0.7426). This suggests that the fecundity of *T. ludeni* on the novel host was already lower than that of *T. urticae* at the onset of the experiment (Fig. 1b, Table 1, electronic supplementary material Table S2-4).

**Table 1.** List of primary antibodies.

|  | Estimate | SE | z value | P value |  |
| --- | --- | --- | --- | --- | --- |
| **The dynamics and performance of the ghost competitor** |  |  |  |  |  |
| (Intercept) (*T. ludeni* on bean) | 2.5867 | 0.1367 | 18.92 | <2e-16 | *** |
| Cucumber | 0.1990 | 0.1965 | 1.01 | 0.3114 |  |
| *T. urticae* | 1.2838 | 0.1720 | 7.46 | 8.47e-14 | *** |
| Cucumber : *T. urticae* | -0.7407 | 0.2468 | -3.00 | 0.0027 | ** |
| **Signature of the ghost competitor on performance of *T. urticae*** |  |  |  |  |  |
| <u>Fecundity assessed on bean</u> |  |  |  |  |  |
| (Intercept) | 3.7098 | 0.0899 | 41.27 | <2e-16 |  |
| <u>Fecundity assessed on cucumber</u> |  |  |  |  |  |
| (Intercept) | 3.2209 | 0.0883 | 36.47 | <2e-16 | *** |
| Initial density *T. ludeni* | 0.0693 | 0.0298 | 2.33 | 0.0199 | * |
| <u>Density during first two months</u> |  |  |  |  |  |
| (Intercept) (*T. ludeni* under comp.) | 1.6678 | 0.3132 | 5.32 | 1.01e-07 | *** |
| *T. urticae* under comp. | -0.1136 | 0.3814 | -0.30 | 0.7658 |  |
| *T. urticae* without comp. | 0.2925 | 0.3744 | 0.78 | 0.4346 |  |
| *T. ludeni* without comp. | 0.2669 | 0.3818 | 0.70 | 0.4845 |  |
| Time | -0.0275 | 0.0099 | -2.79 | 0.0053 | ** |
| *T. urticae* comp. : time | 0.0470 | 0.0117 | 4.00 | 6.26e-05 | *** |
| *T. urticae* no comp. : time | 0.0454 | 0.0114 | 3.97 | 7.35e-05 | *** |
| *T. ludeni* no comp. : time | 0.0351 | 0.0117 | 3.01 | 0.0027 | ** |
| <u>Density during last eight months</u> |  |  |  |  |  |
| (Intercept) (*T. urticae* under comp.) | 2.9066 | 0.0326 | 89.25 | <2e-16 | *** |
| *T. urticae* without comp. | 0.0307 | 0.0462 | 0.67 | 0.5054 |  |
| *T. ludeni* without comp. | -0.4023 | 0.0471 | -8.54 | <2e-16 | *** |
| Time | 2.3787 | 0.5468 | 4.35 | 1.36e-05 | *** |
| Time<sup>2</sup> | -0.9981 | 0.5520 | -1.81 | 0.0706 | . |
| **Performance without interspecific competitor** |  |  |  |  |  |
| (Intercept) *T. urticae* without comp. | 3.5241 | 0.0395 | 89.23 | <2e-16 | *** |
| *T. urticae* under comp. | -0.1119 | 0.0515 | -2.17 | 0.0297 | * |
| *T. urticae* control | -0.1605 | 0.0516 | -3.11 | 0.0019 | ** |

### Signature of the ghost competitor on performance of T. urticae

Throughout the evolutionary experiment, we measured the densities of the populations of both spider mite species. During the first month, the ghost competitor (*T. ludeni*) was still present and the initial density calculated during the first month gave an indication of the pressure exerted by the ghost species on *T. urticae*. We found that the initial density of the ghost competitor positively affected the fecundity of *T. urticae* on the novel host plant (*z* value = 2.33 and *p* = 0.0199). This effect emerged from the start of the experiment and remained stable through time as the minimal model did not include time. Performance on the original host plant, bean, was not related to initial ghost competitor density. Also, the density of *T. urticae* itself or the total initial density was not related to performance on both bean and cucumber (Table 1; electronic supplementary material Table S2-S5).

The initial presence of the ghost competitor influenced the demography of *T. urticae* only slightly. The populations with and without the ghost competitor reached similar equilibrium densities after the ghost competitor went extinct (*t* ratio = -0.666 and *p* = 0.7833), and the increase during the growth phase did not differ significantly (*t* ratio = -0.666 and *p* = 0.7833). At the start no significant difference between densities were found, but after one month the density of *T. urticae* without competition was temporarily significantly higher than the populations of *T. urticae* with *T. ludeni* present (*t* ratio = 3.005 and *p* = 0.0159) (Table 1; electronic supplementary material Table S2-S4).

### Performance without interspecific competitor

We additionally compared the performance of mites from a control population that was maintained on bean plants with mites adapting to cucumber where in both cases the interspecific competitor was replaced by conspecifics. The control population had a significantly lower fecundity on cucumber than the populations grown on cucumber (*t* ratio = -3.110 and *p* = 0.0056), which suggests local adaptation to the novel host plant for the latter group (Fig. 2b; Table 1; electronic supplementary information Table S2-S4).

**Figure 2:**
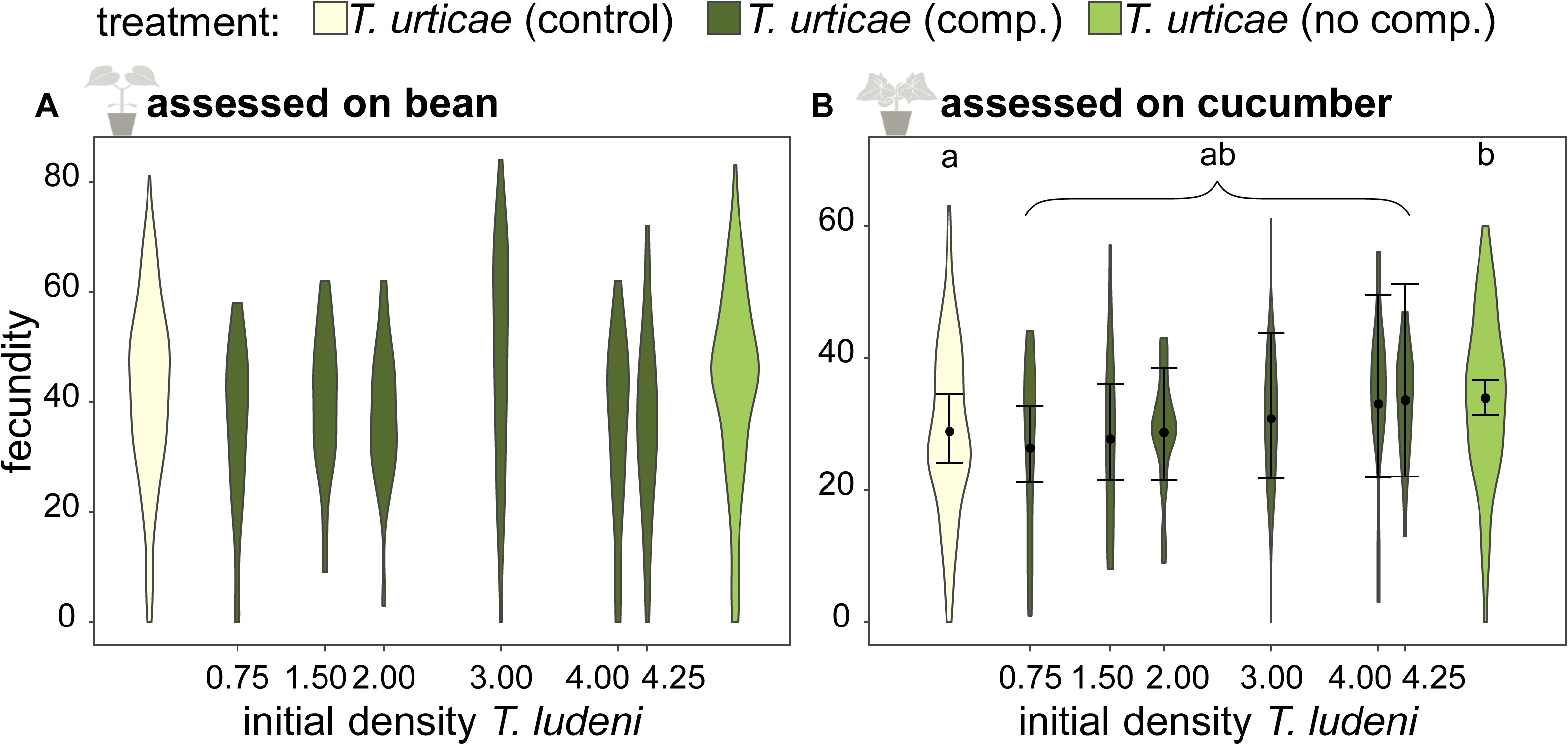
Fecundity affected by ghost competition. On the x-axis the initial density of *T. ludeni* (number of adult females/cm^2^) and the different treatments (*T. urticae* from cucumber but without *T. ludeni*, *T. urticae* with ghost competition of *T. ludeni*, and the control population of *T. urticae* from bean) are presented and on the y-axis the fecundity (number of eggs after six days) of *T. urticae*. This fecundity is averaged over time as this variable was not included in the minimal model. A significant interaction between the density of *T. ludeni* and the fecundity of *T. urticae* was not found when the fecundity was assessed on (A) bean, but it was found when assessed on (B) cucumber. Each violin plot is presenting the observed data, while the points and lines show the means of the model estimate and their 95% confidence interval, respectively.

## DISCUSSION

The process of genetic adaptation to novel environmental conditions is typically studied and understood from the perspective of the available genetic variation and selection pressures as imposed by the environment. Because competing species are an intrinsic part of novel experienced environmental conditions, they are known to mediate sometimes complex evolutionary processes. Here we provide empirical evidence that initial competition between two species can have a long-lasting effect on their performance in a novel environment.

The unintentional rapid extinction of *T. ludeni* seems a logical consequence of the higher attained fecundity of *T. urticae* on the novel host already at the onset of the experiment (Fig. 1b). This higher fecundity and hence higher growth rate increased the chance for better establishment or recovery after disturbance (Turcotte, Reznick & Hare, 2011, 2013). Also, populations from *T. urticae* lived under a higher density than populations from *T. ludeni* (Fig. 1a). The density of *T. urticae* on the measured surface was almost fifty percent more than the density of *T. ludeni* when grown alone. This suggests that *T. urticae* has a higher resource efficiency than *T. ludeni* where the extra energy can be allocated to a higher fecundity. Higher resource efficiency could for instance arise from evolved detoxification mechanisms as often found between herbivores and their hosts (Després, David & Gallet, 2007; Dermauw et al., 2018). After one month the density of populations of *T. urticae* without heterospecific competition was higher than that of populations with heterospecific competition, but this difference vanished together with the extinction of the competitor. This probably means that the ghost competitor decreased the available resources resulting in a lower population size for *T. urticae* (Fig. 1a). We propose that differences in selection pressures exerted by the environment cause the differences within the treatment under ghost competition, but possibly also between this treatment and the populations without competitor.

We have shown that the higher the density of the ghost competitor was, the higher the fecundity of *T. urticae* was on the novel host plant (Fig 2b). We speculate that a higher selection pressure was exerted under a higher initial density of the ghost competitor. This selection pressure eventually led to an increase in fecundity of the focal species. It is known that the competitor, *T. ludeni*, can down-regulate plant defences (Godinho et al. 2016), but this cannot explain the correlation between its higher density and the increased fecundity of the focal species even long after the ghost competitor went extinct, because plants were refreshed weekly.

We found that populations of *T. urticae* without *T. ludeni* reached higher fecundity on the novel host plant than the control population maintained on the initial host plant, implying local adaptation to the novel host (Fig. 2b). This difference in fecundity with the control population was not found for the populations that were initially under competition with *T. ludeni.* A likely explanation for the lack of evidence for adaptation in the treatment under interspecific competition would be the difference in initial population size and subsequent drift effects. Under a scenario of strong drift effects we would expect large differences in performance between replicates; therefore we plotted the fecundity results from the last measured month per replicate (Fig. 3). Since the differences in fecundity assessed on cucumber among replicates are similar with and without *T. ludeni* (and thus with high or low initial population size for *T. urticae*). This indicates that strong drift can be largely discarded as a driver behind the observed evolutionary dynamics.

**Figure 3:**
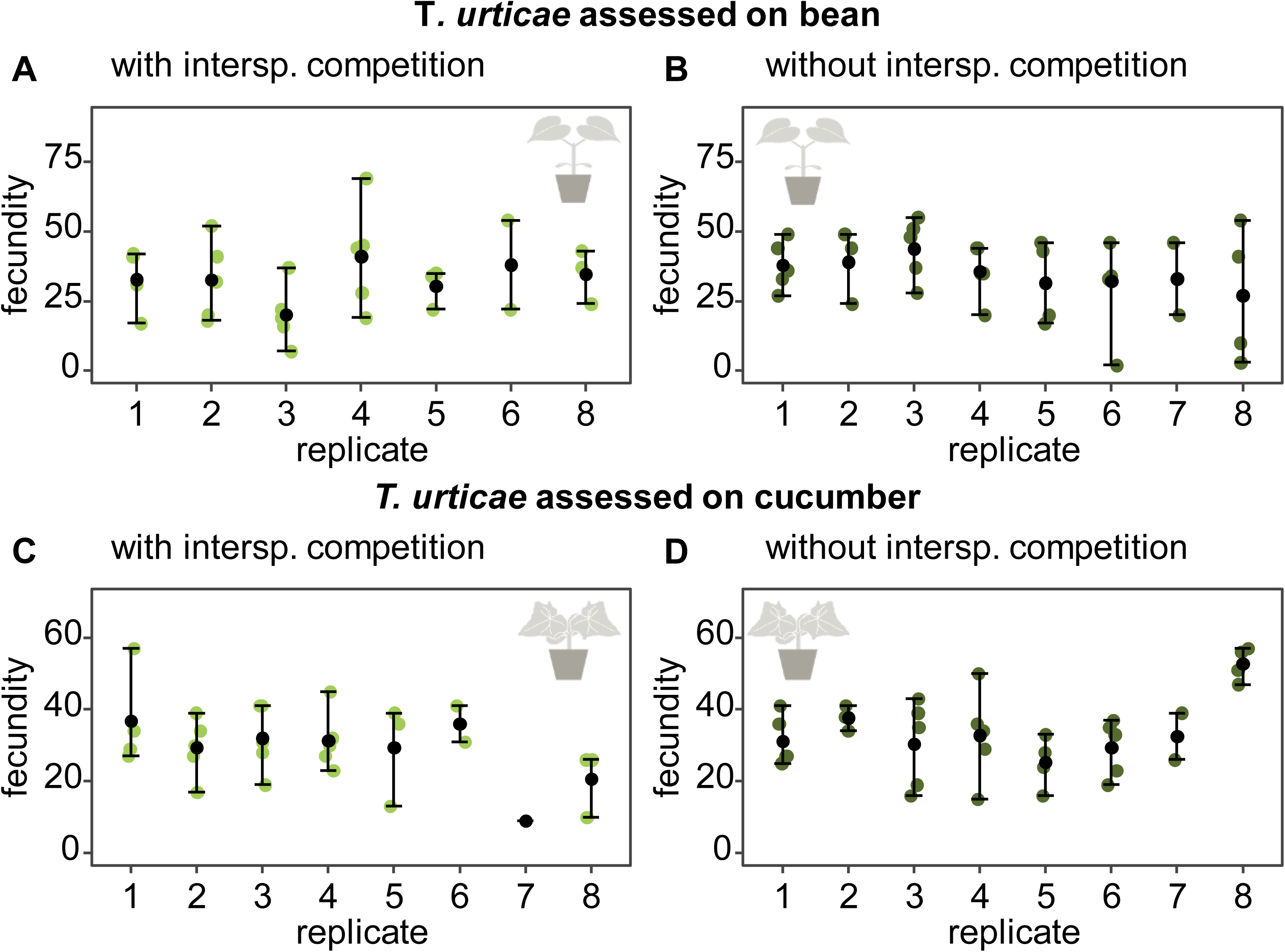
Comparison of fecundity measured after ten months between replicates. The different replicates are given on the x axis and the fecundity (number of eggs after six days) on the y axis. The coloured points are the real measurements, while the black dot and lines present the mean, minimum and maximum value respectively. The upper plots are measurements from *T. urticae* on bean with A) the treatment with *T. ludeni* and B) the treatment without *T. ludeni*. The lower plots are measurements from *T. urticae* on cucumber with C) the treatment with *T. ludeni* and D) the treatment without *T. ludeni*.

The selection pressure of conspecifics was higher than heterospecifics, which may be because an initial larger population sizes, but also due to a larger niche overlap (Bolnick, 2001; Svanbäck & Bolnick, 2007). Furthermore, intraspecific competition is known to push towards ecological specialisation and may thus enhance adaptation to novel host plants (Silvertown, 2004). Intraspecific effects are found to be particularly strong in those cases where communities are affected by indirect interactions such as cascading effects (Des Roches et al., 2018). Many examples are found where indirect interactions are induced after a herbivore attack (e.g. the release of volatiles to attract natural enemies, activation or production of toxins or defensive structures; Zhang *et al*., 2009; Kant *et al*., 2015). Given that we used entire plants, such indirect interactions between herbivores and their host plants likely took place in our experiment.

The history of species in a community can have an impact on interspecific interactions (Fukami, 2015). The magnitudes of such historical contingencies do, however, strongly differ among species and environments (Vannette & Fukami, 2014). Differences in historical contingency have been put forward as an explanation for the fact that some populations can experience radiations, whereas others from the same clade are not capable to achieve this under seemingly similar conditions (Seehausen, 2007). Our results suggest that increased interspecific competition leads to higher selection pressures and thus improved performance (Fig. 2b). Our results coincide in this respect with other empirical work demonstrating that increased competition with heterospecifics increased local adaptation in bunchgrasses (Rice & Knapp, 2008). Similarly, intraguild predation between lizard species increased the selection pressure and led to strong divergence in morphological adaptation as associated with niche specialisation (Stuart et al., 2014).

Nevertheless, we have to be careful with generalising our results. First, we chose small populations sizes as they are more biologically relevant, but this may limit adaptation and establishment (Del Castillo et al., 2011; Yates & Fraser, 2014). We also used populations that have been maintained in the lab for many generations, probably leading to a decrease in genetic variation compared to wild populations. The problems we encountered to create isofemale lines for *T. ludeni* could be an indication of inbreeding depression. However, we are confident that our results are robust as we could still provide evidence for local adaptation in the populations of *T. urticae* without competitor (Fig. 2b). This suggests that the initial amount of genetic variation did not limit *T. urticae* in our study.

Second, it is impossible to add a competitor without changing total population sizes, population densities, or island sizes; all of these are affecting genetic variation and drift (Del Castillo et al., 2011; Alzate, Etienne & Bonte, 2019). As it is known that larger populations usually contain more genetic variation, we chose to standardise this by means of isofemale lines, knowing that this might create differences in drift among treatments. One way to better disentangle the effects of drift from those of selection with our small population size would have been to increase the number of replicates which was difficult for logistical reasons.

Third, our experimental design is not strictly suitable to assess adaptation in the interspecific competition treatment, as we did not keep a control mixed population on bean. Hence, we cannot disentangle the effect of changes in fecundity due to competition (independent of the novel environment) from the effect of competition on adaptation to the novel environment. Although this means that we cannot detect adaptation in the treatment under ghost competition, we did find a positive influence of the density of the ghost competitor on fecundity, meaning that initial selection pressures substantially matter and providing evidence for eco-evolutionary dynamics.

In conclusion, we did find indications for local adaptation in the populations without ghost competition as the performance increased on the novel host compared to a control population. Furthermore, we have shown the importance of initial selection pressures such as ghost competition. Even when one species becomes extinct, the competition signature continues to affect the adaptation process of the successful species. We thus provide experimental evidence on the impact of ghost species on the long-term performance of populations colonizing new environments.

## ACKNOWLEDGEMENTS

We thank Viki Vandomme, Angelica Alcantara, Pieter Vantieghem, Katrien Van Petegem, Stefano Masier, Matti Pisman, Mike Creutz, Hilde De Nil and Johan Bisschop for helping during the research experiments, and to Sarah Magalhães for providing the strains of *T. ludeni.* We thank the Terrestrial Ecology department of Ghent University and the Centre for Ecology, Evolution and Environmental Changes of the University of Lisbon for the spider mite populations. R.S.E. thanks the Netherlands Organisation for Scientific Research (NWO) for financial support through a VICI grant (VICI grant number 865.13.00). K.B. thanks the Special Research Fund (BOF) of Ghent University and the Ubbo Emmius sandwich program of the University of Groningen. DB and RSE are supported by the FWO network EVENET (W0.003.16N) and FWO research grant G018017N.

The authors declare no conflicts of interest.

