## Supplementary material for "Performance in a novel environment subject to ghost competition": Table S1

**Table S1: Overview of the number of samples.**

Summary given per replicate (1 – 8), treatment (*T. urticae* with and without *T. ludeni*, and the control population on bean), and plant species for fecundity assessment (bean, B, and cucumber, C) in time (2 – 10 months) for *T. urticae*. The maximum number per grid is five. Lower numbers indicate adult females that died unnaturally, for instance by drowning before the fecundity assessment. After four months, the cucumber plants did not grow well enough for the fecundity test, therefore we were not able to test fecundity for replicates within that experimental block.

|  |  | 2 months | | 4 months | | 6 months | | 8 months | | 10 months | |
| --- | --- | --- | --- | --- | --- | --- | --- | --- | --- | --- | --- |
|  | replicate | B | C | B | C | B | C | B | C | B | C |
| With *T. ludeni* | 1 | 5 | 5 | 5 |  | 2 | 2 | 5 | 5 | 4 | 4 |
|  | 2 | 5 | 5 | 5 |  | 5 | 5 | 4 | 4 | 5 | 5 |
|  | 3 | 5 | 5 | 4 |  | 5 | 5 | 5 | 5 | 5 | 5 |
|  | 4 | 3 | 3 | 5 |  | 4 | 4 | 5 | 5 | 5 | 5 |
|  | 5 | 5 | 5 | 5 | 4 | 4 | 4 | 4 | 5 | 3 | 3 |
|  | 6 | 5 | 4 | 4 | 4 | 5 | 4 | 5 | 5 | 2 | 2 |
|  | 7 | 5 | 5 | 5 | 5 | 5 | 5 | 4 | 4 | 0 | 1 |
|  | 8 | 3 | 4 | 5 | 4 | 4 | 4 | 5 | 4 | 3 | 3 |
|  | SUM | 36 | 36 | 38 | 17 | 34 | 33 | 37 | 37 | 27 | 28 |
| Without *T. ludeni* | 1 | 4 | 4 | 5 |  | 5 | 5 | 5 | 5 | 5 | 5 |
|  | 2 | 5 | 5 | 4 | 3 | 4 | 4 | 4 | 3 | 3 | 3 |
|  | 3 | 5 | 5 | 5 |  | 3 | 3 | 4 | 4 | 5 | 5 |
|  | 4 | 4 | 4 | 4 |  | 5 | 5 | 5 | 3 | 5 | 5 |
|  | 5 | 4 | 5 | 2 | 2 | 5 | 5 | 5 | 5 | 4 | 4 |
|  | 6 | 5 | 5 | 5 | 5 | 5 | 5 | 4 | 4 | 5 | 5 |
|  | 7 | 5 | 5 | 4 | 4 | 5 | 5 | 4 | 4 | 2 | 2 |
|  | 8 | 5 | 5 | 4 | 4 | 4 | 4 | 5 | 5 | 4 | 4 |
|  | SUM | 37 | 38 | 33 | 18 | 36 | 36 | 36 | 33 | 33 | 33 |
| control | | 31 | 31 | 35 | 17 | 36 | 35 | 36 | 36 | 30 | 31 |
