## Supplementary material for "Performance in a novel environment subject to ghost competition": Table S2

**Table S2: Model selection.**

Overview of the best models based on the lowest AICc with an AICc weight of at least 0.100.

|  | Model | df | LogLik | AICc | Δ AICc | AICc weight |
| --- | --- | --- | --- | --- | --- | --- |
| **The dynamics and performance of the ghost competitor**  maximal model: fecundity ~ plant species * mite species | | | | | | |
| Plant species * mite species | | 5 | - 415.037 | 840.7 | 0.00 | 0.948 |
| **Signature of the ghost competitor on performance of *T. urticae***  **Fecundity assessed on bean -** max. model: fecundity ~ time + initial density *T. urticae* + initial density *T. ludeni* + time : init. dens. *Tu* + time : init. dens. *Tl* + (1\|block/island) | | | | | | |
| No fixed effects | | 4 | -768.410 | 1545.1 | 0.00 | 0.237 |
| Time | | 8 | -764.312 | 1545.5 | 0.45 | 0.190 |
| Initial density *T. urticae* | | 5 | -767.785 | 1545.9 | 0.87 | 0.153 |
| **Fecundity assessed on cucumber -** max. model: fecundity ~ time + initial density *T. urticae* + initial density *T. ludeni* + time : init. dens. *Tu* + time : init. dens. *Tl* + (1\|island) | | | | | | |
| Initial density *T. ludeni* | | 4 | -594.802 | 1197.9 | 0.00 | 0.484 |
| Initial density *T. urticae* + init. dens. *Tl* | | 5 | -594.602 | 1199.6 | 1.74 | 0.203 |
| Initial density *T. urticae* | | 4 | -596.243 | 1200.8 | 2.88 | 0.115 |
| **Demography first two months -** maximal model: density ~ treatment (*T. urticae* with and without competitor, *T. ludeni* with and without competitor) * time + (1\|block/island) | | | | | | |
| Treatment * time | | 11 | -525.602 | 1074.7 | 0.00 | 0.997 |
| **Demography after extinction -** maximal model: density ~ treatment (*T. urticae* with and without competitor, *T. ludeni* without competitor) * time (polynomial) + (1\|block/island) | | | | | | |
| Treatment + time | | 8 | -2888.009 | 5792.2 | 0.00 | 0.91 |
| **Performance without interspecific competitor**  maximal model: fecundity ~ treatment (*T. urticae* control/*T. urticae* with comp./*T. urticae* without comp.) * time + (1\|block/island) | | | | | | |
| Treatment | | 6 | -1846.641 | 3705.5 | 0.00 | 0.751 |
| No fixed effects | | 4 | -1849.890 | 3707.9 | 2.40 | 0.226 |
