## Supplementary material for "Performance in a novel environment subject to ghost competition": Table S3

**Table S3: Chi-square statistics for the maximal models before model selection.** The results for the Wald Chi-square tests are presented for the maximal models.

|  | Independent variables | Chisq | Df | Pr(>Chisq) |  |
| --- | --- | --- | --- | --- | --- |
| **The dynamics and performance of the ghost competitor** | | | | | |
|  | Plant species | 5.1851 | 1 | 0.02278 | * |
|  | Mite species | 56.1019 | 1 | 6.881e-14 | *** |
|  | Plant species : mite species | 9.0064 | 1 | 0.00269 | ** |
| **Signature of the ghost competitor on performance of *T. urticae*** | | | | | |
| Fecundity on bean | Time | 8.4890 | 4 | 0.07522 | . |
|  | Initial density *T. ludeni* | 0.8919 | 1 | 0.34497 |  |
|  | Initial density *T. urticae* | 1.4948 | 1 | 0.22147 |  |
|  | Time : init. dens. *Tl* | 1.3088 | 4 | 0.85987 |  |
|  | Time : init. dens. *Tu* | 4.5696 | 4 | 0.33438 |  |
| Fecundity on cucumber | Time | 4.6468 | 4 | 0.32550 |  |
|  | Initial density *T. ludeni* | 4.1050 | 1 | 0.04276 | * |
|  | Initial density *T. urticae* | 0.6019 | 1 | 0.43786 |  |
|  | Time : init. dens. *Tl* | 1.7385 | 4 | 0.78371 |  |
|  | Time : init. dens. *Tu* | 4.0678 | 4 | 0.39691 |  |
| Demography (first two months) | Treatment | 144.0338 | 3 | <2.2e-16 | *** |
|  | Time | 9.3556 | 1 | 0.00222 | ** |
|  | Time : treatment | 18.6516 | 3 | 0.00032 | *** |
| Demography (after extinction) | Treatment | 104.1507 | 2 | <2.2-16 | *** |
|  | Time (polynomial) | 21.6416 | 2 | 1.998e-05 | *** |
|  | Time (polyn.) : treatment | 3.6009 | 4 | 0.4627 |  |
| **Performance without interspecific competitor** | | | | | |
|  | Treatment | 9.8340 | 2 | 0.007321 | ** |
|  | Time | 0.8189 | 4 | 0.935894 |  |
|  | Treatment : time | 7.9751 | 8 | 0.435902 |  |
