## Supplementary material for "Performance in a novel environment subject to ghost competition": Table S4

**Table S4: Pairwise comparisons adjusted for multiple comparisons (Tukey method).**

| Contrast | Estimate | SE | df | t ratio | p value |  |
| --- | --- | --- | --- | --- | --- | --- |
| **The dynamics and performance of the ghost competitor** | | | | | |  |
| Comparison control population *T. urticae* and *T. ludeni* on bean and cucumber | | | | | |  |
| *T. ludeni* (bean) – *T. ludeni* (cucumber) | -0.199 | 0.197 | 97 | -1.012 | 0.7426 |  |
| *T. ludeni* (bean) – *T. urticae* (bean) | -1.284 | 0.172 | 97 | -7.463 | <0.0001 | *** |
| *T. ludeni* (bean) – *T. urticae* (cucumber) | -0.742 | 0.173 | 97 | -4.279 | 0.0003 | *** |
| *T. ludeni* (cucumber) – *T. urticae* (bean) | -1.085 | 0.176 | 97 | -6.177 | <0.0001 | *** |
| *T. ludeni* (cucumber) – *T. urticae* (cucumber) | -0.543 | 0.177 | 97 | -3.068 | 0.0146 | * |
| *T. urticae* (bean) – *T. urticae* (cucumber) | 0.542 | 0.149 | 97 | 3.629 | 0.0025 | ** |
| **Signature of the ghost competitor on performance of *T. urticae*** | | | | | |  |
| Influence of interspecific competitor on demography (first two months) | | | | | |  |
| Comparison between densities of the different treatments at the start | | | | | | |
| *T. ludeni* (comp.) – *T. urticae* (comp.) | 0.1136 | 0.381 | 181 | 0.298 | 0.9908 |  |
| *T. ludeni* (comp.) – *T. urticae* (no comp.) | -0.2925 | 0.374 | 181 | -0.781 | 0.8629 |  |
| *T. ludeni* (comp.) – *T. ludeni* (no comp.) | -0.2669 | 0.382 | 181 | -0.699 | 0.8973 |  |
| *T. urticae* (comp.) – *T. urticae* (no comp.) | -0.4062 | 0.307 | 181 | -1.325 | 0.5482 |  |
| *T. urticae* (comp.) – *T. ludeni* (no comp.) | -0.3805 | 0.316 | 181 | -1.203 | 0.6256 |  |
| *T. urticae* (no comp.) – *T. ludeni* (no comp.) | 0.0256 | 0.302 | 181 | 0.085 | 0.9998 |  |
| Comparison between densities of the different treatments at the first month | | | | | | |
| *T. ludeni* (comp.) – *T. urticae* (comp.) | -1.2967 | 0.134 | 181 | -9.712 | <0.0001 | *** |
| *T. ludeni* (comp.) – *T. urticae* (no comp.) | -1.6543 | 0.143 | 181 | -11.569 | <0.0001 | *** |
| *T. ludeni* (comp.) – *T. ludeni* (no comp.) | -1.3213 | 0.144 | 181 | -9.147 | <0.0001 | *** |
| *T. urticae* (comp.) – *T. urticae* (no comp.) | -0.3576 | 0.119 | 181 | -3.005 | 0.0159 | * |
| *T. urticae* (comp.) – *T. ludeni* (no comp.) | -0.0247 | 0.120 | 181 | -0.205 | 0.9969 |  |
| *T. urticae* (no comp.) – *T. ludeni* (no comp.) | 0.3329 | 0.117 | 181 | 2.838 | 0.0258 | * |
| Comparison between densities of the different treatments at the second month | | | | | | |
| *T. ludeni* (comp.) – *T. urticae* (comp.) | -2.7070 | 0.372 | 181 | -7.275 | <0.0001 | *** |
| *T. ludeni* (comp.) – *T. urticae* (no comp.) | -3.0160 | 0.370 | 181 | -8.159 | <0.0001 | *** |
| *T. ludeni* (comp.) – *T. ludeni* (no comp.) | -2.3758 | 0.377 | 181 | -6.303 | <0.0001 | *** |
| *T. urticae* (comp.) – *T. urticae* (no comp.) | -0.3090 | 0.261 | 181 | -1.183 | 0.6386 |  |
| *T. urticae* (comp.) – *T. ludeni* (no comp.) | 0.3312 | 0.271 | 181 | 1.224 | 0.6126 |  |
| *T. urticae* (no comp.) – *T. ludeni* (no comp.) | 0.3329 | 0.262 | 181 | 2.445 | 0.0725 | . |
| Interaction between density and time per treatment | | | | | | |
| *T. ludeni* (comp.) | -0.02748 | 0.00985 | 181 | -2.789 | 0.0059 | ** |
| *T. urticae* (comp.) | 0.01953 | 0.00638 | 181 | 3.062 | 0.0025 | ** |
| *T. urticae* (no comp.) | 0.01791 | 0.00582 | 181 | 3.076 | 0.0024 | ** |
| *T. ludeni* (no comp.) | 0.00767 | 0.00629 | 181 | 1.219 | 0.2246 |  |
| Interaction between density and time (comparison of treatments) | | | | | | |
| *T. ludeni* (comp.) – *T. urticae* (comp.) | -0.04701 | 0.01174 | 181 | -4.003 | 0.0005 | *** |
| *T. ludeni* (comp.) – *T. urticae* (no comp.) | -0.04539 | 0.01145 | 181 | -3.965 | 0.0006 | *** |
| *T. ludeni* (comp.) – *T. ludeni* (no comp.) | -0.03515 | 0.01169 | 181 | -3.006 | 0.0159 | * |
| *T. urticae* (comp.) – *T. urticae* (no comp.) | 0.00162 | 0.00863 | 181 | 0.188 | 0.9976 |  |
| *T. urticae* (comp.) – *T. ludeni* (no comp.) | 0.01186 | 0.00896 | 181 | 1.325 | 0.5485 |  |
| *T. urticae* (no comp.) – *T. ludeni* (no comp.) | 0.01024 | 0.00857 | 181 | 1.196 | 0.6304 |  |
| Influence of interspecific competitor on demography (after extinction) | | | | | |  |
| *T. urticae* (comp.) – *T. urticae* (no comp.) | -0.0307 | 0.0462 | 804 | -0.666 | 0.7833 |  |
| *T. urticae* (comp.) – *T. ludeni* (no comp.) | 0.4023 | 0.0471 | 804 | 8.535 | <0.0001 | *** |
| *T. urticae* (no comp.) – *T. ludeni* (no comp.) | 0.4330 | 0.0472 | 804 | 9.168 | <0.0001 | *** |
| **Performance without interspecific competitor** | | | | | |  |
| Investigate local adaptation | | | | | |  |
| *T. urticae* (no comp.) – *T. urticae* (comp.) | 0.1119 | 0.0515 | 453 | 2.174 | 0.0767 | . |
| *T. urticae* (no comp.) – *T. urticae* (control) | 0.1605 | 0.0516 | 453 | 3.110 | 0.0056 | ** |
| *T. urticae* (comp.) – *T. urticae* (control) | 0.0486 | 0.0516 | 453 | 0.941 | 0.6144 |  |
