## Supplementary material for "Performance in a novel environment subject to ghost competition": Table S5

**Model selection and Wald χ² test for the influence of total initial density on fecundity.** Overview of the best models based on the lowest AICc with an AICc weight of at least 0.100.

|  | Model | df | LogLik | AICc | Δ AICc | AICc weight |
| --- | --- | --- | --- | --- | --- | --- |
| **Fecundity assessed on bean -** max. model: fecundity ~ time + total initial density + time : total initial density + (1\|block/island) | | | | | | |
| No fixed effects | | 4 | -768.410 | 1545.1 | 0.00 | 0.293 |
| Total initial density | | 5 | -767.371 | 1545.1 | 0.04 | 0.287 |
| Time | | 8 | -764.312 | 1545.5 | 0.45 | 0.235 |
| Time + total initial density | | 9 | -763.522 | 1546.2 | 1.10 | 0.170 |
| **Fecundity assessed on cucumber -** max. model: fecundity ~ time + total initial density + time : total initial density + (1\|island) | | | | | | |
| No fixed effects | | 3 | -597.445 | 1201.1 | 0.00 | 0.617 |
| Total initial density | | 4 | -597.132 | 1202.5 | 1.48 | 0.294 |

|  | Independent variables | | Chisq | | Df | | Pr(>Chisq) | |
| --- | --- | --- | --- | --- | --- | --- | --- | --- |
| Fecundity on bean | Time | 8.3414 | | 4 | | 0.0798 | | . |
|  | Total initial density | 1.5149 | | 1 | | 0.2184 | |  |
|  | Time : total init. dens. | 4.4325 | | 4 | | 0.3506 | |  |
| Fecundity on cucumber | Time | 4.2049 | | 4 | | 0.3790 | |  |
|  | Total initial density | 0.7819 | | 1 | | 0.3766 | |  |
|  | Time : total init. dens. | 3.8180 | | 4 | | 0.4312 | |  |
